# Differing Associations between Optic Nerve Head Strains and Visual Field Loss in Normal- and High-Tension Glaucoma Subjects

**DOI:** 10.1101/2021.12.15.472712

**Authors:** Thanadet Chuangsuwanich, Tin A. Tun, Fabian A. Braeu, Xiaofei Wang, Zhi Yun Chin, Satish Kumar Panda, Martin Buist, Nicholas Strouthidis, Shamira Perera, Monisha Nongpiur, Tin Aung, Michael JA Girard

**Affiliations:** Ophthalmic Engineering & Innovation Laboratory (OEIL), Singapore Eye Research Institute, Singapore National Eye Center, Singapore; Department of Biomedical Engineering, National University of Singapore, Singapore; Singapore Eye Research Institute, Singapore National Eye Centre, Singapore; Laboratory for Biomechanics and Mechanobiology of Ministry of Education, Beijing Advanced Innovation Center for Biomedical Engineering, School of Biological Science and Medical Engineering, School of Engineering Medicine, Beihang University, Beijing, China; NIHR (National Institute of Health Research) Biomedical Sciences Centre, Moorfields Eye Hospital and UCL Institute of Ophthalmology, London, United Kingdom; Duke-NUS Medical School, Singapore; Institute for Molecular and Clinical Ophthalmology, Basel, Switzerland

**Keywords:** Optic Nerve Head, Normal Tension Glaucoma, Visual Field Loss, Lamina Cribrosa, Ocular Biomechanics

## Abstract

**Purpose:** To study the associations between optic nerve head (ONH) strains under intraocular pressure (IOP) elevation with retinal sensitivity in glaucoma subjects.

**Design:** Clinic based cross-sectional study.

**Participants:** 229 subjects with primary open angle glaucoma (subdivided into 115 high tension glaucoma (HTG) subjects and 114 normal tension glaucoma (NTG) subjects).

**Methods:** For one eye of each subject, we imaged the ONH using spectral-domain optical coherence tomography (OCT) under the following conditions: **(1)** primary gaze and **(2)** primary gaze with acute IOP elevation (to approximately 33 mmHg) achieved through ophthalmodynamometry. A 3-dimensional (3D) strain-mapping algorithm was applied to quantify IOP-induced ONH tissue strain (i.e. deformation) in each ONH. Strains in the pre-lamina tissue (PLT)and the retina, the choroid, the sclera and the lamina cribrosa (LC) were associated (using linear regression) with measures of retinal sensitivity from the 24-2 Humphrey visual field test (Carl Zeiss Meditec, Dublin, CA, USA). This was done globally, then locally according to the regionalization scheme of Garway-Heath et al.

**Main Outcome Measures:** Associations between ONH strains and values of retinal sensitivity from visual field testing.

**Results:** For HTG subjects, we found that **(1)** there were significant negative linear associations between ONH strains and retinal sensitivity (p<0.001) (on average, a 1% increase in ONH strains corresponded to a decrease in retinal sensitivity of 1.1 dB), **(2)** high strain regions co-localized with anatomically-mapped regions of high visual field loss, **(3)** the strongest negative associations were observed in the superior region and in the PLT. In contrast, for NTG subjects, no significant associations between strains and retinal sensitivity were observed except in the supero-temporal region of the LC.

**Conclusion:** We found significant negative associations between IOP-induced ONH strains and retinal sensitivity in a relatively large glaucoma cohort. Specifically, HTG subjects who experienced higher ONH strains were more likely to exhibit lower retinal sensitivities. Interestingly, this trend was in general less pronounced in NTG subjects, which could suggest a distinct pathophysiology between the two glaucoma subtypes.

## Introduction

Loss of vision in glaucoma is preceded by a cascade of cellular events leading to damage of the retinal ganglion cell (RGC) axons. From a biomechanical perspective, the neural and connective tissues around the lamina cribrosa (LC) experience intraocular pressure (IOP)-related stresses and strains; these could in turn elicit RGC damage (directly or indirectly) at the level of the LC.^1^ Apart from IOP, the biomechanical stresses in the LC can also be influenced by age,^2^ the presence of ocular comorbidities (e.g. myopia)^3^ and a fluctuating cerebrospinal fluid pressure (CSFP).^4^ Increased biomechanical stresses due to the aforementioned factors (and their interactions) could ultimately influence an already fragile ONH,^5^ and the resulting deformations could correlate with functional loss in glaucoma. Using OCT imaging and computational algorithms, several authors have successfully reported IOP-induced biomechanical deformations within the ONH tissues in terms of ‘strains’,^6–11^ which can in-turn be used for calculations of tissue stiffness/compliance.^12^ Apart from our earlier study^13^ which reported that the amount of ONH strains alleviated due to trabeculectomy was associated with the extent of visual field loss based on a few glaucoma subjects, no studies have investigated associations between local ONH strains and visual field loss in glaucoma. We believe that such associations would provide strong evidence to support the clinical usefulness of ONH strains extracted from OCT images for both glaucoma diagnosis and prognosis.

In addition, with the high prevalence of normal tension glaucoma (NTG), especially in Asian populations,^14, 15^ we believe that it would be valuable to compare the aforementioned associations between NTG and high-tension glaucoma (HTG) subjects. The differences in relationships between IOP-induced strain and functional loss in glaucoma in NTG and HTG subjects can possibly highlight differences in etiologies, if they exist, between the two glaucoma subgroups.

The aim of this study was to investigate the relationship between *in vivo* local ONH strains (induced by IOP elevation) and retinal sensitivity values measured from 24-2 Humphrey visual field tests in NTG and HTG subjects. To this end, we applied a strain/displacement mapping algorithm, based on our previous works,^6, 7, 9, 16^ to optical coherence tomography (OCT) images to quantify *in vivo* ONH strains in each subject. We then utilized a map derived by Garway-Heath et al.^17^ to co-localize anatomical ONH regions to their corresponding visual field counterparts.

## Methods

Our goal was to quantitatively map 3D ONH strains in NTG and HTG subjects under IOP elevation and correlate the ONH strains to the retinal sensitivity values obtained from the 24-2 Humphrey visual field test. To this end, we first imaged each subject’s ONH in baseline gaze using OCT, and subsequently, under acute IOP elevation. ONH tissue deformations were mapped using a digital volume correlation (DVC) algorithm applied to the pair of OCT volumes. Such deformations were then statistically correlated to retinal sensitivity values using linear regressions across the entire ONH tissues and across regions of the visual field using a spatial map provided by Garway Heath et al.^17^ Below is a detailed description of our methodology.

### Subjects Recruitment

We recruited 229 subjects with primary open angle glaucoma (115 subjects with HTG and 114 with NTG) from glaucoma clinics at the Singapore National Eye Centre. We included subjects aged more than 50 years old, of Chinese ethnicity (predominant in Singapore), with a refractive error of ±3 diopters, and excluded subjects who underwent prior intraocular/orbital/brain surgeries, subjects with past history of strabismus, ocular trauma, ocular motor palsies, orbital/brain tumors; with clinically abnormal saccadic or pursuit eye movements; subjects with poor LC visibility in OCT (<50% en-face visibility); subjects with known carotid or peripheral vascular disease; or with any other abnormal ocular conditions. Glaucoma was defined as glaucomatous optic neuropathy, characterized as loss of neuroretinal rim with vertical cup-to-disc ratio >0.7 or focal notching with nerve fiber layer defect attributable to glaucoma (based on a clinical observation) and/or asymmetry of cup- to disc ratio between eyes >0.2, with repeatable glaucomatous visual field defects (independent of the IOP value) in at least 1 eye. Glaucomatous visual field defect was defined the following conditions were present: (1) glaucoma hemifield test outside normal limits, (2) a cluster of ≥ 3, non-edge, contiguous points on the pattern deviation plot, not crossing the horizontal meridian with a probability of < 5% being present in age-matched normal (one of which was < 1%) and (3) Pattern Standard Deviation (PSD) < 0.05; these were repeatable on two separate occasions. We used gonioscopy to determine the angle status of the recruited patients - the eye was defined as closed if the posterior trabecular meshwork could not be seen in more than one quadrant by dark room gonioscopy in primary gaze position without indentation. NTG subjects had consistently low/normal IOP (=<21 mmHg) before treatment in the study eye; HTG subjects had elevated IOP (>21 mmHg) before treatment in the study eye.

Each subject underwent the following ocular examinations: (1) measurement of refraction using an autokeratometer (RK-5; Canon, Tokyo, Japan); (2) measurement of axial length, central corneal thickness (CCT) and anterior chamber depth (ACD) using a commercial device (Lenstar LS 900; Haag-Streit AG, Switzerland). For each tested eye we performed a visual field test using a standard achromatic perimetry with the Humphrey Field Analyser (Carl Zeiss Meditec, Dublin, CA).

This study was approved by the SingHealth Centralized Institutional Review Board and adhered to the tenets of the Declaration of Helsinki. Written informed consent was obtained from each subject.

### Visual Field Testing

Subjects with visual field deficits related to diabetic retinopathy or any other optic neuropathies, or advanced glaucoma (MD of less than −20 dB) precluding acute elevation of IOP were excluded. Unreliable visual field test results with a false-positive error > 15%, or a fixation loss > 33% were excluded.^18^ Thresholding Algorithm (Swedish Interactive Testing Algorithm) standard 24-2 program were selected. For every selected subject the retinal sensitivity map from the Humphrey Visual Field report was used in our study.

### OCT Imaging

We selected one eye at random for each subject and we imaged the ONH with spectral-domain OCT (Spectralis; Heidelberg Engineering GmbH, Heidelberg, Germany). The imaging protocol was similar to that from our previous work.^6^ In brief, we conducted a raster scan of the ONH (covering a rectangular region of 15° x 10° centered at the ONH), comprising of 97 serial B-scans, with each B-scan comprising of 384 A-scans (**Figure 1a**). The average distance between B-scans was 35.1 μm and the axial and lateral B-scan pixel resolution were on average 3.87 μm and 11.5 μm respectively. All B-scans were averaged 20 times during acquisition to reduce speckle noise. Each eye was scanned two times under two conditions – baseline OCT position, and acute IOP elevation. Each subject was administered with 1.0% Tropicamide to dilate the pupils before imaging.

**Figure 1.**
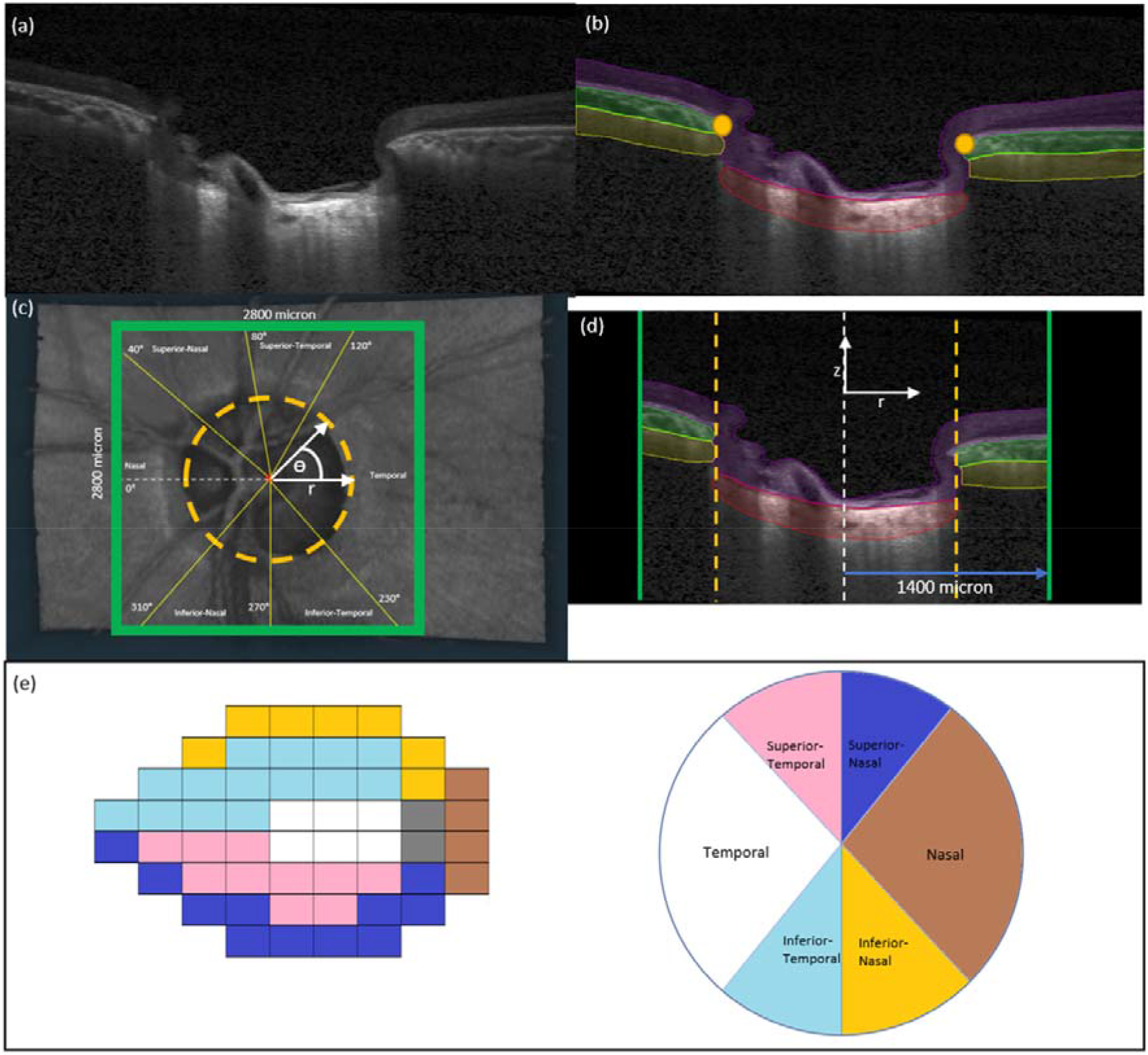
**(a)** A single B-scan obtained from the OCT machine without any image enhancement **(b)** Automatic segmentation of the B-scan in (a). Four tissues were segmented – Pre-Lamina tissue (purple), Choroid (green), Sclera (yellow) and LC (red) In addition, BMOs (orange dots) were automatically marked for each B-scan **(c)** Anterior-surface view of the ONH. The ONH center (white star) was identified from the best-fit circle to the BMOs (orange-dotted line). Green square defines our region of interest to be cropped from the OCT volume with 2800µm length on each side. Yellow lines divide the ONH into 6 based on the angular degree from the reference white-dotted line. White axes represent a cylindrical coordinate system (□ is circumferential direction, r is radial direction). **(d)** A B-scan view after we apply cropping to the OCT volumes. Black region was not considered for our analysis. The length from central line (white-dotted line) to the cropping border (green line) is 1400 µm. White axes represent a cylindrical coordinate system (z is axial direction, r is radial direction). **(e)** Regionalization of the 24-2 visual field test (left) to the ONH tissue (right) according to the Garway-Heath scheme.

### OCT imaging during acute IOP elevation

For each eye in baseline position, we applied a constant force of 0.65 N to the temporal side of the lower eyelid using an opthalmodynamometer, as per a well-established protocol.^6, 7^ The force was applied for approximately 2 to 3 minutes throughout the OCT scan duration and varied for no more than one minute across subject. This force raised IOP to about 35 mmHg and was maintained constant. IOP was then re-assessed with a Tono-Pen (Reichert Instruments GmbH, Munich, Germany), and the ONH was imaged with OCT in baseline position immediately (within 30 seconds) after the IOP was measured.

### ONH Reconstruction through Automatic Segmentation

For each ONH, we automatically segmented the following tissue groups - the PLT+retina, the choroid, the visible portion (excluding posterior limit) of the peripapillary sclera, and the visible portion (excluding posterior limit) of the LC (**Figure 1b**). This was done using a deep-learning algorithm (a dilated-residual U-Net) based on our previous work.^19^ The quality of the segmentation was rigorously validated based on a large dataset and human experts.^19^ Here, segmentation was required so as to summarize strains for any chosen tissue group in subsequent steps. BMO points were also automatically extracted with a custom algorithm. Note that BMO points lie within a plane (the BMO plane), and such a plane can be used as a horizontal reference plane for each ONH.

### ONH Regional Division

To ensure un-biased comparisons between groups (NTG vs HTG) for 3D deformations/strains, we first limited our en-face field-of-view to a region of 2800 × 2800 µm^2^ centered on the BMO center for all ONHs (**Figure 1c**). Each ONH was further divided into six regions (nasal, supero-nasal, supero-temporal, temporal, infero-temporal and infero-nasal) according to the regionalization scheme of Garway-Heath et al.^17^ (**Figure 1c**). To aid in calculation of strains, we defined a cylindrical coordinate system with the BMO center as the origin and the axes are defined as the radial distance (r), the circumferential angle (□), and the axial distance (z; depth) according to **Figure 1c-d**.

Each region of the ONH was spatially associated to the 24-2 visual field test points according to the Garway-Heath scheme shown in **Figure 1e**.

### In Vivo Displacement and Strain Mapping of the ONH

We used a commercial DVC module (Amira, 2020.3, Waltham, Massachussets: Thermo Fisher Scientific) to map the three-dimensional deformations between the baseline scan and acute IOP elevation scan for each patient. The working principle of this commercial DVC module is similar to our prior DVC implementation^9^, albeit with an improved speed efficiency. Details of the DVC algorithm used in this study is provided in **Appendix A**. Briefly, each ONH morphology was sub-divided into ∼4,000 cubic elements, and ∼3,500 nodes (points), at which locations 3D displacements (vectors) were mapped following an acute IOP elevation. We then derived the **(1)** effective strain, **(2)** the radial strain Err, **(3)** the circumferential strain E□□, **(4)** the axial strain Ezz and **(5)** the shear components Er□, Erz, and E□z, **(6)** the maximum principal strain E1 (corresponding to the maximum local tensile strain), and the **(7)** minimum principal strain E3 (corresponding to the maximum local compressive strain) from the 3D displacements. The directions for the local coordinate system (r, □ and z) are shown in **Figure 1**. The effective strain is a convenient local measure of 3D deformation that takes into account both the compressive and tensile effects. In other words, the higher the compressive or tensile strain, the higher the effective strain.^6, 9, 16^ Circumferential strain takes into account the circular geometry of the eye and it highlights the effect of the circumferential forces (i.e., hoop stress) borne by the scleral canal under IOP elevation.^20^ Axial strain corresponds to the axial direction of the eye and it highlights the ONH tissue compression under IOP elevation. Details of the strain derivation is provided in **Appendix B**, and further validation of the DVC and its effects on strain in **Appendix C**.

### Statistical Analysis

To study the associations between IOP-induced strain and retinal sensitivity, we calculated the mean values of ONH effective strain (all tissues and all regions) and the mean values of retinal sensitivity for each subject. We then fit a linear regression model using MATLAB for each subject group with the effective strain as a dependent variable and the retinal sensitivity as an independent variable. We also performed similar analyses separately for the PLT+Retina, Choroid, Sclera and LC.

To study the regional association of ONH effective strain and retinal sensitivity, we calculated the mean values of effective strain in each region and their respective mean values of retinal sensitivity for each subject. In essence, each subject had 6 data points (corresponding to mean effective strain and mean retinal sensitivity in each region) and we fit a linear regression model on each subject. To represent an average regional association across all subjects, we also fit a linear regression model on all data points across every subject of each group.

To investigate the association of each tissue, each region, and each subject group to each type of strain (effective strain, circumferential strain, and axial strain), we performed in total 142 linear fits (number of combinations of the aforementioned parameters). We performed a Bonferroni correction to adjust the statistically significant p-value to the number of comparisons and reported significant associations based on the adjusted p-value.

Statistical significance level was set at 0.05 for all linear model fits. We compared the strength of the linear relationships based on the estimated coefficients.

## Results

### Demographics and IOP elevation

A total of 229 Chinese subjects were recruited (consisting of 115 subjects with HTG and 114 with NTG). We excluded 24 HTG subjects and 23 NTG subjects from the study due to a low LC visibility (<50% of the BMO area on either the baseline or elevated IOP scan) and/or unreliable visual field results. LC visibility was determined based on the top-view area that the segmented LC occupied with respect to the BMO area, as was performed in our previous study.^7^ Therefore, 91 HTG subjects and 91 NTG subjects were included in the final analysis. Out of 91 HTG subjects, 32 subjects were female. Out of 91 NTG subjects, 41 subjects were female.

There were no significant differences (p>0.05) across both groups in terms of age [HTG: 69 ± 5, NTG: 67 ± 6], systolic blood pressure [HTG: 141 ± 16 mmHg, NTG: 140 ± 20 mmHg], diastolic blood pressure [HTG: 75 ± 9 mmHg, NTG: 74 ± 9 mmHg], axial length [HTG: 24.2 ± 1.0 mm, NTG: 24.4 ± 1.0 mm], visual field mean deviation [HTG: −7.54 ± 5.05 dB, NTG: −6.56 ± 4.91 dB], pattern standard deviation [HTG: 7.18 ± 3.79 dB, NTG: 7.22 ± 3.05 dB], baseline IOP (on the day of the experiment) [HTG: 17.3 ± 2.9 mmHg, NTG: 16.0 ± 2.5 mmHg], IOP during ophthalmodynamometer indentation [HTG: 34.5 ± 7.0 mmHg, NTG: 34.8 ± 6.5 mmHg], BMO area [HTG: 2.45 ± 0.63 mm^2^, NTG: 2.37 ± 0.58 mm^2^], and the mean ΔIOP (IOP with ODM application minus baseline IOP) [HTG: 17.4 mmHg ± 7 mmHg, NTG: 18.5 mmHg ± 7 mmHg].

The average IOP at diagnosis for was 25.2 ± 3.5 mmHg for HTG subjects and 17.2 ± 2.0 mmHg for NTG subjects.

In terms of visual field, there were no significant differences (p>0.05) across both groups in terms of MD values [HTG: −7.46 ± 5.9 dB, NTG: −6.17 ± 5.8 dB], mean retinal sensitivity values [HTG: 21.54 ± 4.6 dB, NTG: 19.43 ± 9.40 dB] and mean pattern deviation values [HTG: −5.51 ± 5.61 dB, NTG: −5.13 ± 5.63 dB].

Almost all subjects were treated with prostaglandin analogue, except for 4 subjects in the NTG group. For those who received prostaglandin analogue, the number of drops taken by HTG patients was higher (1.85 ± 0.83 drops/day) as compared to NTG patients (1.38 ± 0.64 drops/day). However, we found no significant correlation between ONH effective strain and the number of eyedrops taken (p = 0.2).

### Significant Negative Associations between Effective Strain and Retinal Sensitivity in HTG Subjects

Across the entire ONH (all tissues, all regions), we found a significant negative association between the average retinal sensitivity (dB) and the average effective strain for HTG subjects (p < 0.001) with a regression coefficient (β) of −1.1 (**Figure 2a**). In other words, a 1% increase in effective strain correlate to 1.1 dB decrease in retinal sensitivity.

**Figure 2.**
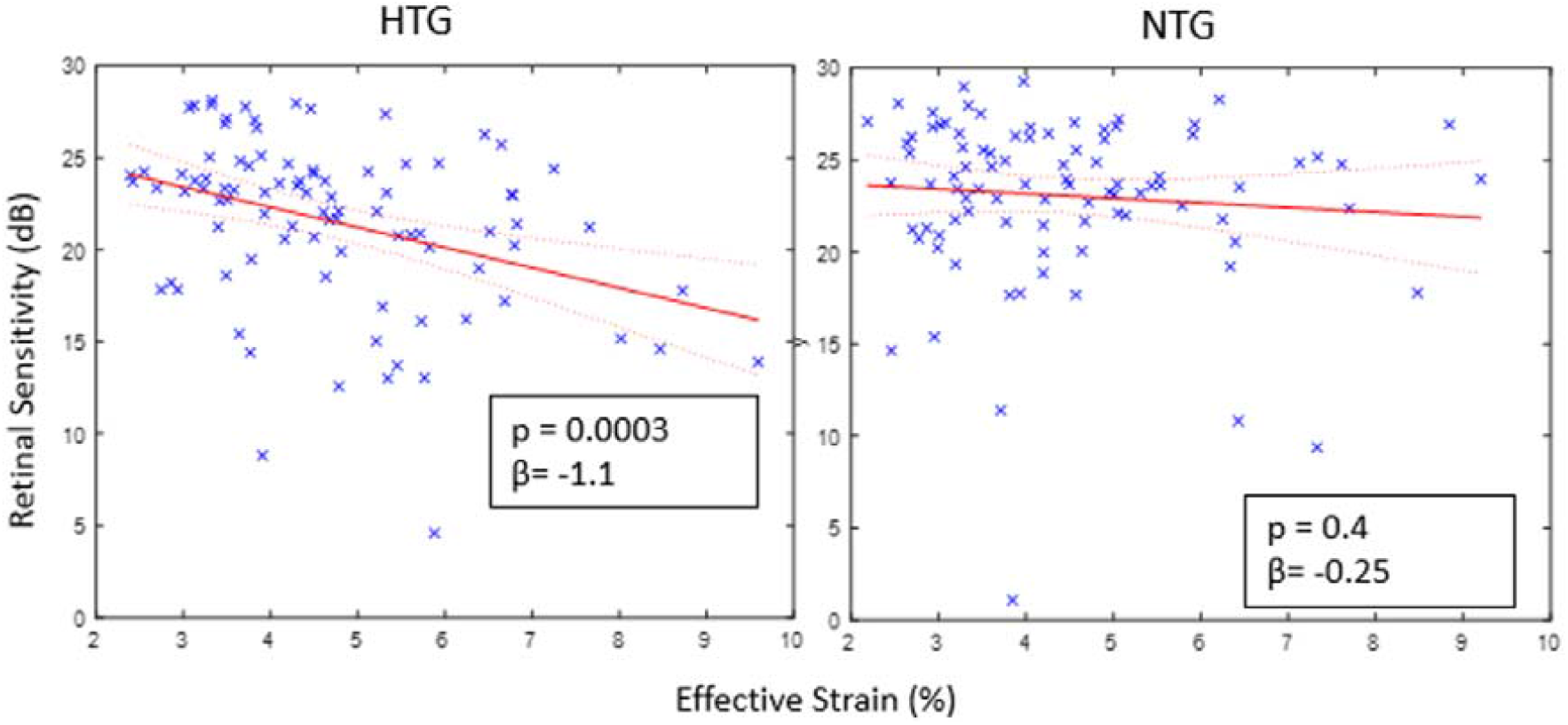
A scatter plot of retinal sensitivity (dB) against effective strain (%) with regression lines (red) and confidence bounds (red-dotted) for HTG and NTG subjects. p represents p-value of the linear fit and β represents the coefficient of the fit.

For NTG subjects, we found no significant association (p = 0.4) between the average retinal sensitivity (dB) and the average effective strain (**Figure 2b**).

### Strongest Negative Association in the PLT Tissue of HTG subjects

For all ONH tissues (LC, PLT, Choroid and sclera – **Figure 3a-d**), we found significant negative associations between the average retinal sensitivity (dB) and the average effective strain for HTG subjects (all tissues p-values <0.001). The tissues exhibiting the strongest negative associations for HTG subjects were the PLT (β = −1.1), followed by the sclera (β = −1.0), the LC (β = −0.9) and the choroid (β = −0.9).

**Figure 3.**
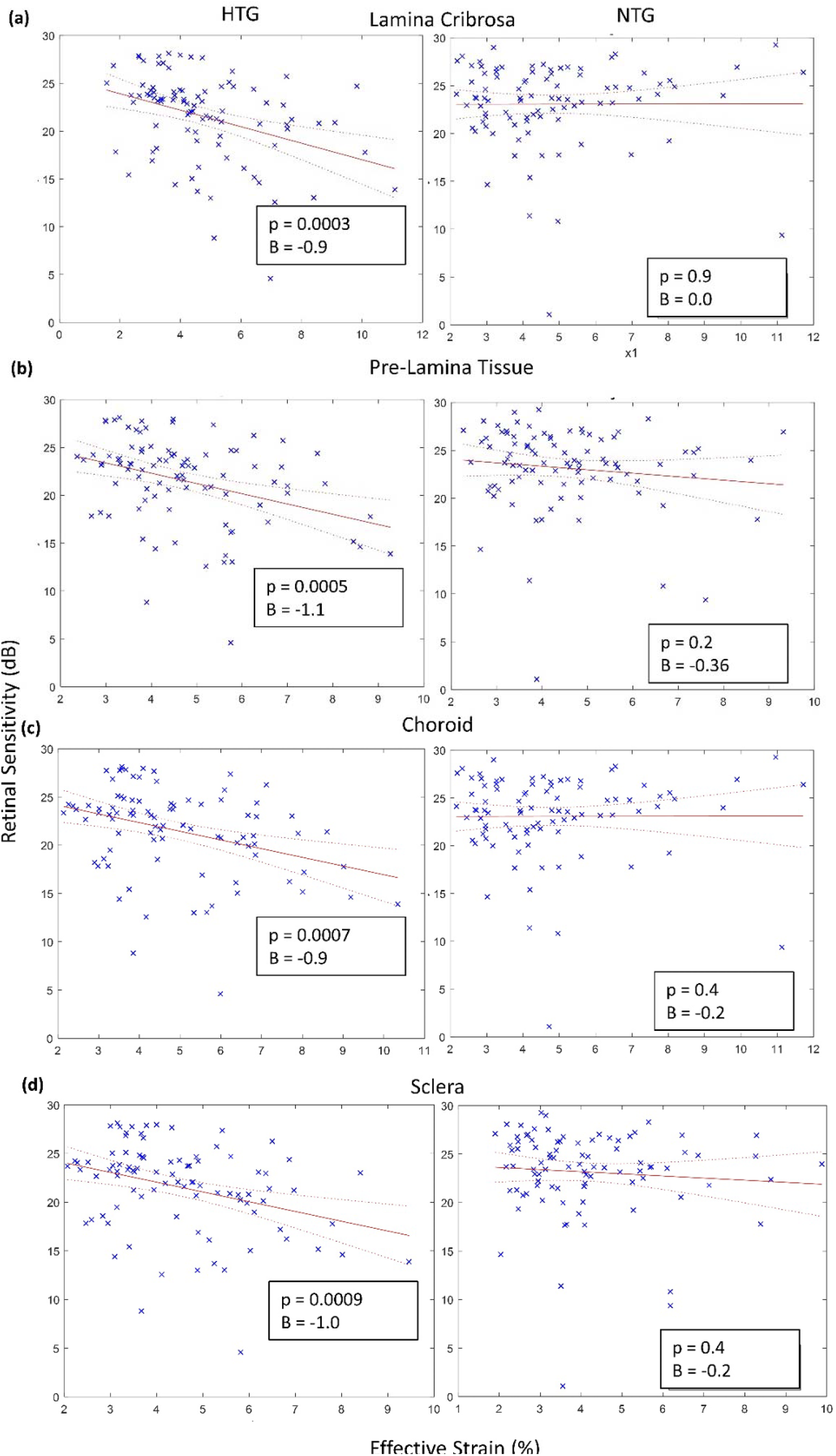
A scatter plot of retinal sensitivity (dB) against effective strain (%) with regression lines (red) and confidence bounds (red-dotted) for the **(a)** lamina cribrosa **(b)** pre-lamina tissue **(c)** choroid and **(d)** sclera of the HTG and NTG subjects. p represents p-value of the linear fit and β represents the coefficient of the fit.

For NTG subjects, there were no significant associations between the average retinal sensitivity (dB) and the average effective strain for all tissues (p > 0.05, **Figure 3a-d**).

### ONH Regions with High Effective Strains Corresponds to Regions with Low Retinal Sensitivity

Across all HTG and NTG subjects, there is a significant association between regions with high strains and regions with low retinal sensitivities (p < 0.001 and p = 0.004 respectively, **Figure 4a**).

**Figure 4.**
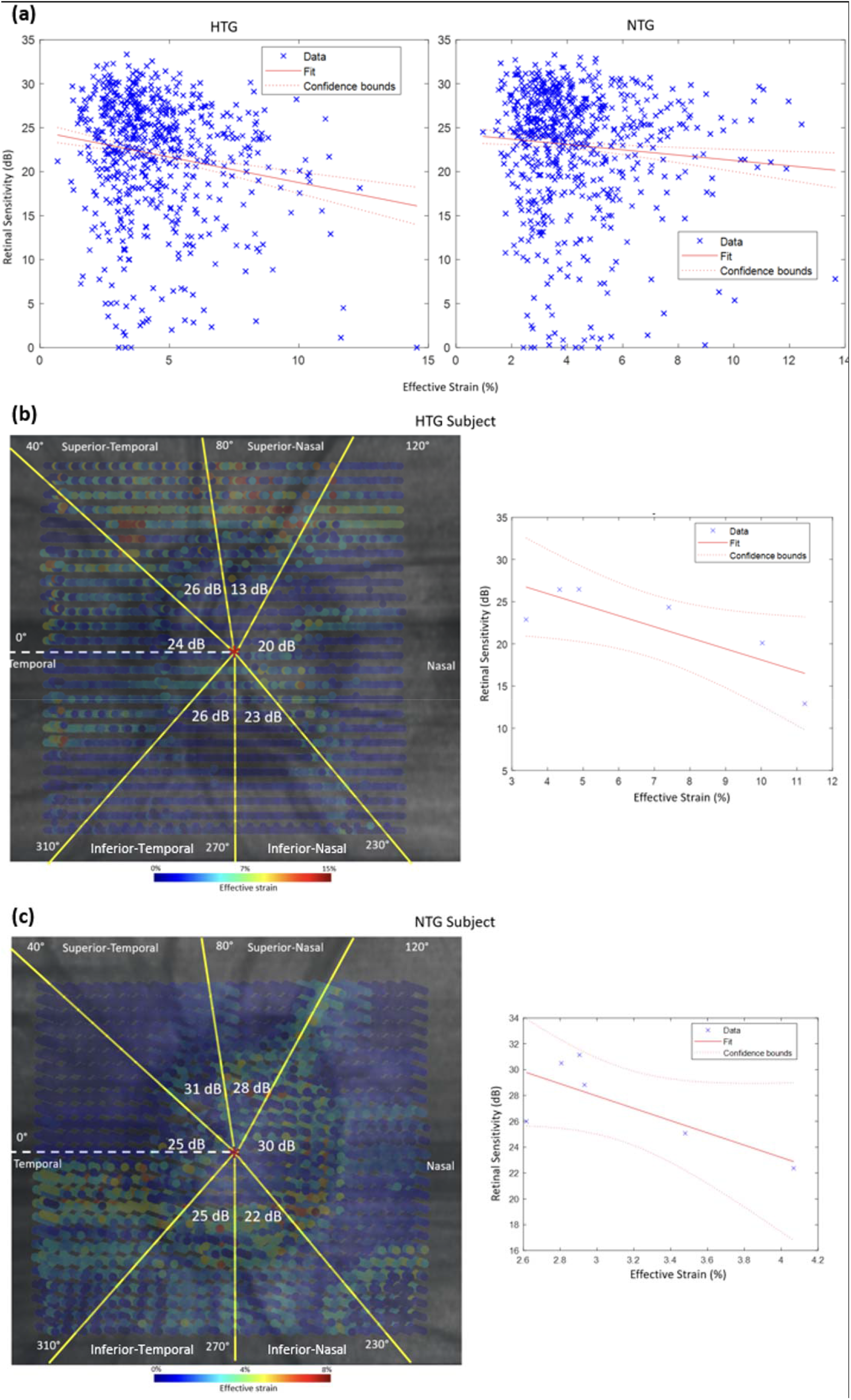
**(a)** Scatter plots of all subjects (6 data points for each subject) of the average retinal sensitivity values against the effective strain. **(b)** Overlay of calculated effective strain on the en-face image of a single HTG subject’s ONH (left) with corresponding colour bar to represent the strain values. Average retinal sensitivity values for each region are given in white text. Scatter plot (right) with linear regression line showing linear associations between average effective strain and average retinal sensitivity in each region. **(c)** Same representation as (a) for an NTG subject.

To illustrate this association, **Figure 4b** shows a co-localization of the highest average strain value (11.3%) and the lowest average retinal sensitivity value (13 dB) in the superior-nasal region of an HTG subject. Likewise, **Figure 4c** shows a co-localization of the highest average strain value (4.1%) and the lowest average retinal sensitivity value (22 dB) in the inferior-temporal region of an NTG subject.

### Ranking of all Associations

In our analysis, we found that the most important (and significant after a Bonferroni correction) strain measures (with respect to all strain measures) were circumferential strain and the effective strain (**Table 1**). We have provided additional details of top 20 associations between each strain and the retinal sensitivity and also summarized the state of strains (tensile vs compressive) in **Appendix D**.

**Table 1.**
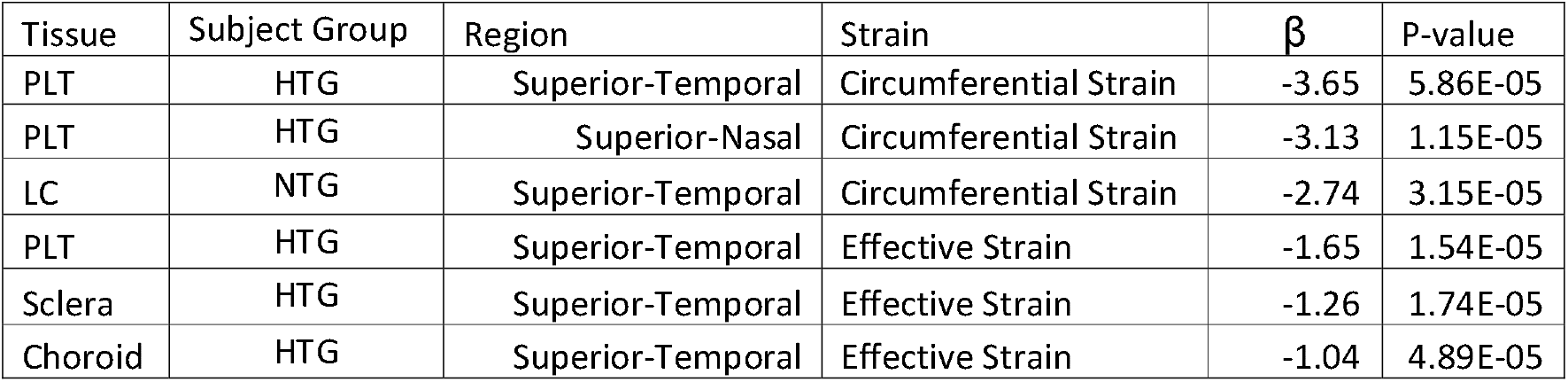
Ranking of all significant associations (Bonferroni corrected from 142 total associations) between each type of strain and retinal sensitivity values with respect toe each tissue type and each region.

| Tissue | Subject Group | Region | Strain | $\beta$ | P-value |
| --- | --- | --- | --- | --- | --- |
| PLT | HTG | Superior-Temporal | Circumferential Strain | -3.65 | 5.86E-05 |
| PLT | HTG | Superior-Nasal | Circumferential Strain | -3.13 | 1.15E-05 |
| LC | NTG | Superior-Temporal | Circumferential Strain | -2.74 | 3.15E-05 |
| PLT | HTG | Superior-Temporal | Effective Strain | -1.65 | 1.54E-05 |
| Sclera | HTG | Superior-Temporal | Effective Strain | -1.26 | 1.74E-05 |
| Choroid | HTG | Superior-Temporal | Effective Strain | -1.04 | 4.89E-05 |

All significant associations were negative (i.e., higher strains correspond to lower retinal sensitivities). The strongest association for HTG subjects was observed in the superior-temporal region of the PLT tissue with circumferential strain as a measure, with a 1% increase in circumferential strain corresponding to approximately 3.65 dB reduction in retinal sensitivity.

We found a single significant association for NTG subjects, specifically in the superior-temporal region of the LC tissue with circumferential strain as a measure, with a 1% increase in circumferential strain corresponding to approximately 2.74 dB reduction in retinal sensitivity.

All significant associations were observed in the superior region, with 5 of them occurring in the supero-temporal region and one in the supero-nasal region.

## Discussion

In this study, we investigated the associations between IOP-induced ONH strain and retinal sensitivity, with respect to each ONH tissue and each ONH region. We also compared the associations across two group of glaucoma subjects (NTG and HTG). For HTG subjects, we found significant negative associations between ONH strains induced by IOP elevation and retinal sensitivity, whereas for NTG subjects, such associations were less significant.

We found that IOP-induced ONH effective strain had a significant association with retinal sensitivity in HTG subjects (**Figure 2**). This association has been reported before in our earlier study,^16^ however, the key differences were that **(1)** we reported associations with respect to IOP lowering (from trabeculectomy), whereas herein, with respect to IOP elevation (through ophthalmo-dynamometry); **(2)** our population was significantly larger (229 glaucoma subjects herein vs 9 glaucoma subjects in our previous study). In addition, we found that the association was negative, and on average an increase of 1% in ONH effective strains corresponded to a decrease of 1.1 dB in retinal sensitivity. Interestingly, significant negative associations were found for any given tissue, whether neural or connective (**Figure 3**). According to the biomechanical theory of glaucoma,^21^ IOP-induced strain could potentially: (1) directly induce mechanical damage to the RGCs,^22^ (2) disrupt the microcapillary blood flow at the LC, choroid and the retina^23–25^ and (3) disrupt the axoplasmic flow.^26^ Therefore, it would be plausible that eyes with structurally weaker ONHs (and thus higher strains) would be more susceptible to visual field loss. The results from this study suggest that a simple mechanical test (IOP elevation via ophthalmodynamometer) could provide insights into the structural-functional relationship for glaucoma subjects. Further studies should be carried out to investigate whether such relationships could be used to predict glaucoma progression.

Of note, we observed no global associations between ONH strains and retinal sensitivity in NTG subjects (**Figure 2**) on average for all ONH tissues nor for each specific ONH tissue. This is an interesting finding since all HTG and NTG subjects were matched demographically and both groups had similar visual field indices. On the surface, this could imply that IOP elevation (or its fluctuation) may not influence ONH biomechanics in NTG subjects in a way that directly translates to visual field loss. It may support the notion of a different aetiology for NTG subjects that is IOP independent.^27, 28^ For instance, several factors have been proposed to explain the development and progression of glaucoma at lower levels of IOP, such as a vascular deficiency,^27^ a low CSFP,^29, 30^ structural weaknesses of ocular tissues,^31, 32^ or the presence of a strong optic nerve head traction force during eye movements.^6, 33–36^ It is also plausible that the stress-strain curve of NTG subjects was starkly different to that of the HTG subjects, due to the differences in ocular biomechanical properties,^37^ and that our controlled force application elicited the strain response in the region of the curve that happened to be non-associative to the visual field.

In terms of regional associations, we observed that regions of high strains were co-localized with anatomically-mapped regions of high visual field loss for both groups of subjects, with a stronger association observed in HTG subjects (**Figure 4a**). In glaucoma, the superior and inferior region of the optic nerve are more susceptible to ganglion cell loss,^38^ which gives rise to what is described as an hourglass-like pattern of neuronal death. This follows the anatomical feature of the supportive collagenous beams at the LC, where the beams are less dense (resulting in larger optic nerve passages) in the superior and inferior regions as compared to the nasal and temporal region.^39^ It is likely that the superior and inferior regions of the LC are more susceptible to IOP-related forces (resulting in higher deformation and strains) due to the lesser presence of the supporting collagenous beams. In addition, the remodelling process of the connective tissue during glaucoma development may also contribute to the focal damage to the nerve cells, as this process could change the LC pores diameters, the collagen beam thickness and other LC morphologies, all of which influence the LC responses’ to IOP elevation.^40^ In our study, we observed that the superior regions (both superior-nasal and superior-temporal) of the ONH had the strongest associations (**Table 1**). Although our study agrees with the aforementioned focal pattern neuronal loss in the superior regions, we did not find significant association in the inferior regions, which are also the common site for ONH damage in glaucoma.^41–43^ This discrepancy may be attributed to the different stages of glaucoma disease as remodelling of the extracellular matrix can lead to a rapid change of ONH tissues responses to a mechanical stress. Our subjects consist of a broad spectrum of glaucoma severity (from mild to severe) and it is possible that the inferior-temporal structural weaknesses may be more prominent in a high severity glaucoma. Of note, we observed a significant association in NTG subjects in the superior-temporal region of the LC tissue (Table 1).

In terms of strains measures, circumferential strains provided the strongest associations (**Table 1**). Circumferential ONH strains can primarily be attributed to IOP-related stresses acting on the peripapillary sclera,^21^ this latter being supported by a circumferentially-arranged network of collagen and elastin fibres.^44^ Circumferential ONH strains can be thought of as local measures of scleral canal expansion,^45^ where higher circumferential strains may imply that the ONH’s ability to withstand IOP-related forces was compromised, possibly from the weakening of the collagenous ring surrounding the optic disc.^46^ We also found that the average circumferential strain was tensile for both PLT and LC tissues (**Appendix D)**. The circumferential strain represents the ‘hoop’ stress and the tensile value indicates an ‘expansionary’ force under IOP elevation; this observation is in line with other studies.^20, 21^ Therefore, it would be sensible for circumferential strains to be strongly associated with visual field loss as observed in this study. Additionally, we observed that the ONH tissues were under compressive axial strains, which was particularly high in the LC tissue; this observation is also in line with other studies.^47, 48^ Tissue-wise, the PLT provided the strongest association. The significant thinning of the PLT tissue in glaucoma disease, resulting from the loss of a wide variety of cells within the retina, including ganglion cells, muller cells and others,^49^ may compromise the ability of the PLT to tolerate IOP-induced forces causing an increase (that corresponds to the severity of glaucoma disease) in IOP-induced strains in the PLT.

We would like to emphasize that our analyses reported herein (both global and regional associations) were based on the ‘raw’ retinal sensitivity values. Since we controlled for ethnicity, age, other eye diseases and past ocular surgeries, we decided to utilize the raw sensitivity values as they represented an unprocessed functional data. The average raw sensitivity values were also used in our previous paper^16^ and others^50, 51^ which study the structure-function relationship within the same demographics. We also performed a global analysis using on MD values and a regional analysis using clusters of TD values and reached the same conclusions as stated in this manuscript – **(1)** strong correlation between visual field for and strains for HTG subjects but not for NTG subjects and **(2)** strong correlations between regions of high strains and regions of high visual field defect. However, for PSD and PD values, their correlations to strains became less significant across both subject groups, which were expected because PSD and PD values account for localized deviations in visual field patterns with respect to the normal population, whereas our strain is an absolute measure without any adjustments to ‘normal strains’ patterns. We have presented the additional analysis in **Appendix G**.

Several limitations in this study warrant further investigation. First, we did not categorize glaucoma into different stages. Since different stages of glaucoma could have distinct biomechanical manifestations,^52–54^ our analysis would miss out on those differences and instead represent an average trend of all glaucoma stages.

Second, our definition of NTG vs HTG (based on an IOP value of 21 mmHg), introduces an additional degree of uncertainties with respect to our analysis for those subjects with IOP values of ∼21 mmHg before treatment. A binary classification of subjects would also exclude the possibility to investigate the threshold IOP as a continuous variable. However, this definition is still the standard for classifying NTG patients and it allows our results to be compared to pre-existing literatures.^55–57^ We have performed additional analysis to treat the threshold IOP as a continuous variable by varying the threshold IOP from 18 to 23 mmHg and found that our conclusion (significant correlations between visual field and strains in HTG subjects but not in NTG subjects) was still the same from an IOP threshold of 19 to 23 mmHg. The detail of this additional analysis is provided in **Appendix E**.

Third, the actual level of IOP raised via ODM varied both inter and intra subjects based on several factors: (1) the ocular pulse,^58^ (2) the viscoelastic behaviour of the ONH connective tissue,^59^ which may cause the level of IOP raised to fall slightly after the initial compression as observed in a study by Kazemi et al.^60^ and, (3) to a lesser extent, the response of the eyelid tissue to the ODM. In our current study, we observed a relatively high standard deviation of 7 mmHg for the level of IOP raised for both groups of subjects. In this study, we have performed an additional analysis to adjust our effective strains according to the level of IOP raised (that was patient specific) and found that our conclusion still held true. We have included this additional analysis in **Appendix F**.

Fourth, during the experiment, we used Tonopen to measure the baseline and elevated IOP, due to its compact dimension which does not interfere with the OCT imaging. Several studies^61^ have shown that IOP values measured from the Tonopen may be inaccurate in for high values of IOP (IOP > 35 mmHg) as compared to applanation-based tonometer.

Fifth, errors in both displacement and strain could occasionally occur. The errors observed here could arise from various sources such as OCT registration errors (intrinsic to the OCT machine), rotation of subjects’ head during OCT acquisition, OCT speckle noise and IOP fluctuations from ocular pulsations,^62^ all of which were difficult to control. However, displacement error magnitudes were still lower than the OCT voxel resolution (Appendix C); thus, these errors should not have significantly affected the observed trends.

Sixth, the Garway-Heath map was created based on European subjects which, may not translate to our subjects. Also, the number of visual field points assigned to each region based on the Garway-Heath map were unbalanced (13 visual field points correspond to the inferior-temporal region as opposed to only 4 points corresponding to the nasal region), which could introduce ascertainment bias regionally. In addition, the papillomacular bundle which corresponds to the central visual field defect, may enter the optic disc closer to the infero-temporal zone,^63^ but the Garway-Heath does not subdivide the temporal regions into superotemporal and inferotemporal quadrants. To complement the structure-function analysis using Garway-Heath map, we also compared the regional RNFL thickness to the strains in each regional of the ONH. Interestingly, we still found significant negative correlations between RNFL thickness and the corresponding strains in each region (Superior, Inferior, Nasal and Temporal) for HTG subjects but not for NTG subjects. The detail for this additional analysis is provided in **Appendix H**.

Lastly, our results strongly suggest that ‘effective strain’ is an important measure to link with visual field loss (**Table 1** and **Appendix D**), and it should be a strong candidate for clinical applications of biomechanics in glaucoma. The effective strain has several advantages: **(1)** it summarizes the multidimensional state of strain into a single value, taking into account compression, tension, and shear, making it easy to interpret clinically; **(2)** it is indepedent of the coordinate system employed; **(3)** it is essentially a von Mises strain, which is commonly used in mechanical engineering for damage analysis and is thus very relevant for glaucoma, and **(4)** it has been employed by another groups^64^ in ONH tracking studies.

In conclusion, we found that there were significant negative associations between IOP-induced ONH strains and retinal sensitivity. In short, HTG subjects who experienced higher strains due to acute IOP elevation also exhibited lower retinal sensitives. We also identified the tissue (PLT) and the ONH regions (Superior regions) where these associations were the strongest. In addition, weaker associations were observed in NTG subjects, which could be indicative of differing aetiologies between the NTG and HTG groups.

## Acknowledgements

Acknowledgement is made to **(1)** the donors of the National Glaucoma Research, a program of the BrightFocus Foundation, for support of this research (G2021010S [MG]), **(2)** the Singapore Ministry of Education, Academic Research Funds, Tier 2 (R-397-000-280-112; R-397-000-308-112 [MG]) & Tier 1 (R-397-000-294-114 [MG]), **(3)** the “Retinal Analytics through Machine learning aiding Physics (RAMP)“ project supported by the National Research Foundation, Prime Minister’s Office, Singapore under its Intra-Create Thematic Grant “Intersection Of Engineering And Health” - NRF2019-THE002-0006 awarded to the Singapore MIT Alliance for Research and Technology (SMART) Centre, (4) SingHealth DukeNUS, Academic medicine research grant (AM/SU053/2021 [TAT]) and (5) the National Natural Science Foundation of China (12002025 [XW]).

## Abbreviations/Acronyms

IOP: Intraocular Pressure
CSFP: Cerebrospinal Fluid Pressure
ONH: Optic Nerve Head
LC: Lamina Cribrosa
RGC: Retinal Ganglion Cell
NTG: Normal-Tension Glaucoma
HTG: High-Tension Glaucoma
OCT: Optical Coherence Tomography
DVC: Digital Volume Correlation
PLT: Pre-Lamina Tissue
BMO: Bruch’s Membrane Opening

## Notes

**Financial Support:** This work was supported by (1) the Singapore Ministry of Education, Academic Research Funds, Tier 2 (R-397-000-280-112; R-397-000-308-112) (2) Singapore Ministry of Education, Academic Research Funds, Tier 1 (R-397-000-294-114) (3) the National Medical Research Council (Grant NMRC/STAR/0023/2014) (4) SingHealth DukeNUS, Academic medicine research grant (AM/SU053/2021) and (5) National Natural Science Foundation of China (12002025).

**Conflict of Interest:** All authors declare no conflict of interest except Michael J.A. Girard (Abyss Processing Pte Ltd).

### Competing Interest Statement

The authors have declared no competing interest.

