## Supplemental Appendix for "Differing Associations between Optic Nerve Head Strains and Visual Field Loss in Normal- and High-Tension Glaucoma Subjects"

A. **Details of DVC algorithm**

We used a commercial software module (Amira *XDigitalVolumeCorrelation* Extension) to map the three-dimensional deformation of the following OCT volume pairs – (1) baseline OCT vs acute IOP elevation (2) baseline OCT vs adduction, and (3) baseline OCT vs abduction – for each patient. The software utilized a robust Finite Element (FE)-based DVC approach which is often called a ‘global’ approach.^24, 25^ Unlike the ‘local’ DVC approach used in our previous works,^26^ a FE-based DVC requires the FE mesh of the entire region of interest as a method of discretization, in contrast to a ‘sub-volumes’ region of interest for the local approach. Therefore, in the global approach, the whole volume of interest is considered in each iteration and the method used by Amira *XDigitalVolumeCorrelation* was described in detail by Hild et al.^24^ Briefly, FE-based DVC problem consisted of solving the following equation –

$$f\left( \boldsymbol{x} \right)=g\left( \boldsymbol{x}+u\left( \boldsymbol{x} \right) \right) Eq. 1$$

where $x$ is a position of any voxel, $f$ is reference volume voxel intensity (in grayscale), $g$ is deformed volume voxel intensity (in grayscale) and $u$ is the unknown displacement field. Solving the above equation is equivalent to minimizing the sum of squared differences of the correlation residual $\Phi_{c}(x)$ –

$$\Phi_{c}\left( \boldsymbol{x} \right)=|f\left( \boldsymbol{x} \right)-g\left( \boldsymbol{x}+u\left( \boldsymbol{x} \right) \right)| Eq. 2$$

A C8 finite element (and its associated trilinear shape function) was used to discretize the problem^25^ and to generate the following linear system to be solved –

$$\left[ \boldsymbol{M} \right]\left\{ \delta u \right\}=\left\{ b \right\}Eq. 3$$

where $\left[ M \right]$ is the DVC matrix, $\{\delta u\}$ is the correction vector to vector $u$ at each iteration and $\{b\}$ is the DVC vector that needs to be minimized.

The algorithm converges when $\delta u$ is smaller than 10^-3^ micrometer. To summarize, *XDigitalVolumeCorrelation* needs three main inputs – (1) hexahedron mesh of the ROI (2) convergence criterion which is the tolerance ($\delta u$), and (3) a baseline image volume with its corresponding deformed image volume. In this study, we generated a hexahedron mesh using AMIRA software mesh generator by specifying a mesh size of 100x100x100 micrometer or approximately 9x3x26 voxels in X, Y and Z directions for each volume. This resulted in a hexahedron mesh with approximately 3500 nodes per volume. The DVC algorithm calculated the displacements of each cube’s vertices (the cube’s corners). We set the tolerance of $\delta u$ to be 1x10^-4^ for each volume.

B. **Derivation of strains**

From the displacement field, we calculated a deformation gradient tensor (***F***) and we derived a Green Lagrange strain tensor consisting of 6 components according to the following equation –

$$E = \frac{1}{2}\left( F^{T}\times F-I \right)$$

where E is Green Lagrange strain tensor and I is identity matrix. In this study, we reported effective strain as the main measure of strain at a tissue node. We calculated effective strain from the principal components (E_1_, E_2_, E_3_) of the Green Lagrange strain tensor (E) according to the following equation –

$$E_{eff} =\sqrt{\frac{{(E_{1}-E_{2})}^{2}+{(E_{1}-E_{3})}^{2}+{(E_{2}-E_{3})}^{2}}{2}}$$

where $E_{eff}$ is the effective strain. Note that effective strain is a single positive index that represents both compressive and tensile effects. In other words, higher compressive strain or tensile strain will result in a higher effective strain.

C. **Validation of DVC method**

The DVC algorithm baseline displacement and strain error were estimated from the following artificial cases (performed on a baseline volume of healthy subjects) – (1) 20-micrometer rigid body translation along a positive x-direction, (2) a 2-degree clockwise rotation about the center of the ONH, (3) 4% tension, (4) 4% compression along the x-direction, (5) radial expansion from the geometric center of the LC with a mean effective strain of approximately 4%, (6) 3% compression along the z-direction, (7) a combination of case 5 and 6 and (8) case 6 with an addition of gaussian noise (mean 0 and standard deviation of 0.05). We also estimated the baseline error due to variation in subject’s body position and motion between each scan by comparing repeated baselines (N = 3) scans from a single subject. We quantify the errors in terms of the following parameters: displacement magnitudes in the X, Y and Z direction and effective strains.

For case (1), the average error in displacements were 0.086 ± 0.10 micron in the x-direction, 0.028 ± 0.27 micron in the y-direction, 0.035 ± 0.33 micron in the z-direction and the average error in effective strain value was 0.0024 ± 0.001%. For case (2), the average error in displacements were 0.024 ± 0.66 micron in the x-direction, 0.033 ± 0.43 micron in the y-direction, 0.017 ± 0.068 micron in the z-direction and the average error in effective strain value was 0.0070 ± 0.005%. For case (3), the average error in displacements were 0.26 ± 0.65 micron in the x-direction, 0.002 ± 0.016 micron in the y-direction, 0.015 ± 0.19 micron in the z-direction and the average error in effective strain value was 0.0003 ± 0.002%. For case (4), the average error in displacements were 0.36 ± 0.71 micron in the x-direction, 0.031 ± 0.019 micron in the y-direction, 0.011 ± 0.22 micron in the z-direction and the average errors in effective strain value was 0.0025 ± 0.002%. %. For case (5), the average error in displacements were 0.44 ± 0.76 micron in the x-direction, 0.042 ± 0.033 micron in the y-direction, 0.018 ± 0.31 micron in the z-direction and the average errors in effective strain value was 0.0065 ± 0.003%. %. For case (6), the average error in displacements were 0.015 ± 0.22 micron in the x-direction, 0.011 ± 0.018 micron in the y-direction, 0.033± 0.35 micron in the z-direction and the average errors in effective strain value was 0.007 ± 0.003%. %. For case (7), the average error in displacements were 0.51 ± 0.31 micron in the x-direction, 0.032 ± 0.018 micron in the y-direction, 0.035± 0.35 micron in the z-direction and the average errors in effective strain value was 0.0074 ± 0.003%. For case (8), the average error in displacements were 0.65 ± 0.37 micron in the x-direction, 0.035 ± 0.022 micron in the y-direction, 0.038± 0.37 micron in the z-direction and the average errors in effective strain value was 0.0077 ± 0.003%.

Overall, for the artificial deformation cases, the maximum error in displacements along the X, Y and Z direction was less than 5% of the voxel resolution. The maximum error in effective strain was 0.5%.

For three repeated baseline scans of a healthy subject (N1, N2, N3), the average displacement errors for the pair N1-N2 were 3.2 ± 3.8 micron in the x-direction, 1.2 ± 1.9 micron in the y-direction, 3.3 ± 5.0 micron in the z-direction and the average error in effective strain was 1.0 ± 0.1%. For the pair N1-N3, the average displacement errors were 1.9± 3.1 micron in the x-direction, 1.5 ± 2.8 micron in the y-direction, 5.3 ± 4.8 micron in the z-direction and the average error in effective strain was 0.9 ± 0.08%. For the pair N2-N3, the average displacement errors were 3.4± 3.2 micron in the x-direction, 1.5 ± 2.9 micron in the y-direction, 5.5 ± 4.3 micron in the z-direction and the average error in effective strain was 1.1 ± 0.08%. Overall, for the repeated baseline scans of a single subject, the maximum error in displacements was approximately 30% of the voxel resolution and the maximum error in effective strain was 1.2%. Additionally, the average error for each strain component based on a repeated baseline scan was as follow: the absolute error for Err was 0.0098, Eɵɵ 0.0009, Ezz 0.0012, Erɵ 0.0008, Erz 0.001, and Eɵz 0.0008%.

D. **Expanded Ranking of Correlations and Regional Strain Summary**

We have provided additional ranking of correlations based on p-values according to the table below.

| Tissue | Cond | Region | Strain | Gradient | R2 | P-value | Ranking |
| --- | --- | --- | --- | --- | --- | --- | --- |
| PLT | HTG | Superior-Nasal | Circumferential Strain | -3.13595 | 0.195358 | 1.15E-05* | 1 |
| PLT | HTG | Superior-Temporal | Effective Strain | -1.65957 | 0.190355 | 1.54E-05* | 2 |
| Sclera | HTG | Superior-Temporal | Effective Strain | -1.26537 | 0.188233 | 1.74E-05* | 3 |
| LC | NTG | Superior-Temporal | Circumferential Strain | -2.74768 | 0.256306 | 3.15E-05* | 4 |
| Choroid | HTG | Superior-Temporal | Effective Strain | -1.04907 | 0.173559 | 4.89E-05* | 5 |
| PLT | HTG | Superior-Temporal | Circumferential Strain | -3.65601 | 0.166757 | 5.86E-05* | 6 |
| PLT | HTG | Superior-Temporal | Maximum Principal Strain | -2.07307 | 0.143162 | 0.000217 | 7 |
| Sclera | HTG | Nasal | Maximum Principal Strain | -1.84088 | 0.197708 | 0.000331 | 8 |
| Sclera | HTG | Nasal | Effective Strain | -0.86175 | 0.134911 | 0.000342 | 9 |
| PLT | HTG | Nasal | Minimum Principal Strain | -1.85854 | 0.132412 | 0.000392 | 10 |
| Choroid | HTG | Superior-Temporal | Circumferential Strain | -2.54913 | 0.135774 | 0.000412 | 11 |
| Choroid | HTG | Nasal | Effective Strain | -0.79791 | 0.131341 | 0.000415 | 12 |
| Choroid | HTG | Nasal | Maximum Principal Strain | -1.56173 | 0.13142 | 0.000446 | 13 |
| Choroid | HTG | Nasal | Minimum Principal Strain | -1.4436 | 0.126358 | 0.000586 | 14 |
| PLT | HTG | Nasal | Effective Strain | -0.88315 | 0.118329 | 0.000842 | 15 |
| PLT | HTG | Inferior-Temporal | Axial Strain | -4.07798 | 0.11779 | 0.000867 | 16 |
| LC | HTG | Temporal | Radial Strain | -3.45257 | 0.127035 | 0.000876 | 17 |
| Choroid | HTG | Superior-Nasal | Circumferential Strain | -1.6352 | 0.117158 | 0.001174 | 18 |
| LC | HTG | Nasal | Erz | -3.85504 | 0.118395 | 0.001262 | 19 |
| PLT | HTG | Superior-Nasal | Minimum Principal Strain | -1.86436 | 0.105504 | 0.001682 | 20 |

Table A1: Top 20 associations based on the p-value of linear regressions based on each type of strain, tissue and region with respect to the average retinal sensitivity. * denotes significant correlations after bonferroni adjustments.

Additionally, to further investigate the state of strains (tensile/compressive), we have plotted out the average values of the Err, Eɵɵ and Ezz in each region of the PLT and LC tissue in **Figure A1**. We found that Eɵɵ is tensile for both PLT and LC tissues. Eɵɵ represents the ‘hoop stress’ and the positive Eɵɵ indicates an ‘expansionary’ force under IOP elevation; this observation is in line with other studies.^[1],^^[2]^ In addition, we observed that the tissues were under compressive axial strains, which is particularly high in the LC tissue; this observation is also in line with other studies.^[3],^^[4]^

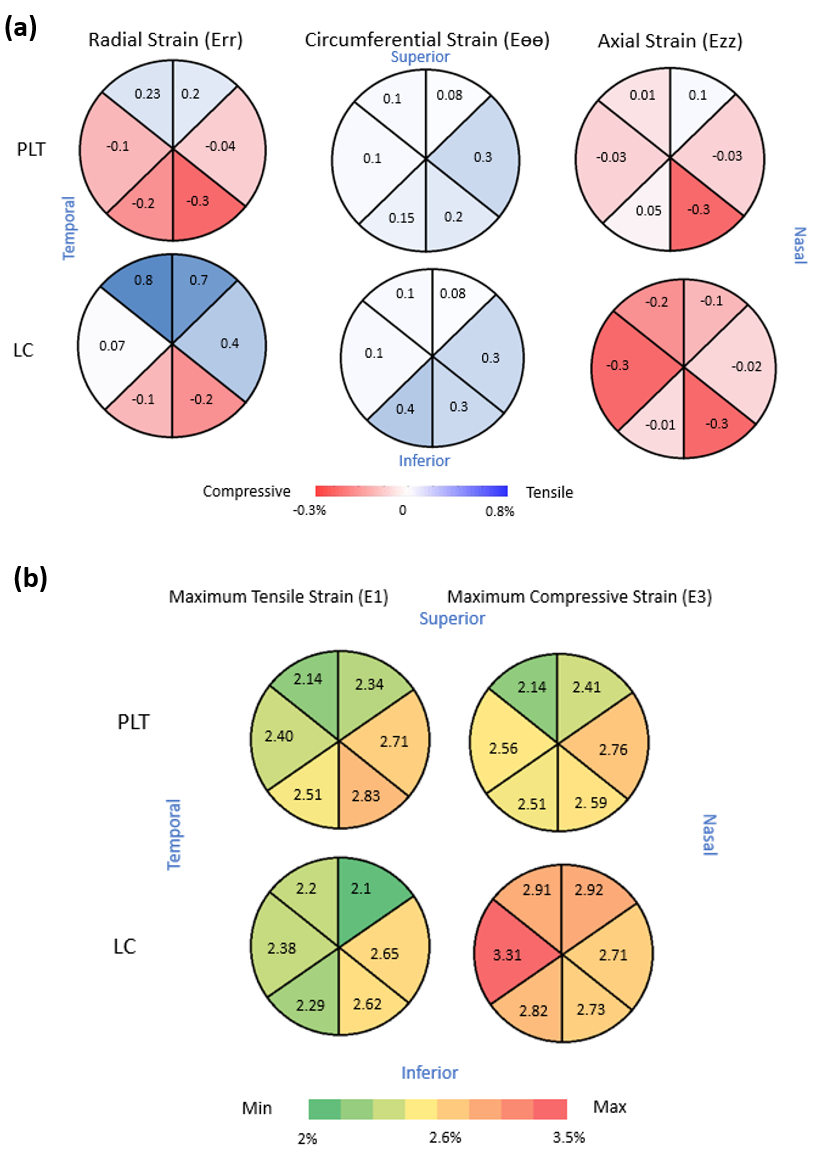

Figure A1: (a) Average radial strain, circumferential strain and axial strain with respect to each region in the PLT and LC tissue. (b) Average E1 and E3 with respect to each region in the PLT and LC tissue.

E. **Effects of Varying IOP Threshold on the Global Correlations**

We can treat the threshold IOP for binarizing the subjects as a pseudo-continuous variable by varying the threshold of ‘normal’ IOP from 18 to 23 mmHg in an increment of 1 mmHg (i.e., our lowest threshold of 18 mmHg will classify NTG as those with <= 18 mmHg and HTG otherwise). We then performed linear regression between retinal sensitivity and IOP induced strains based on our set of thresholds, and we presented our findings in the table below.

| IOP threshold (NTG <= threshold, HTG otherwise) | Linear regression between LC strains and HTG subjects (p-value and beta coefficient) | Linear regression between LC strains and NTG subjects (p-value and beta coefficient) |
| --- | --- | --- |
| 18 | p-value: 0.16, beta: -0.27 | p-value: 0.6, beta: -0.30 |
| 19 | p-value: 0.049, beta: -0.43 | p-value: 0.8, beta: -0.09 |
| 20 | p-value: 0.04, beta: -0.53 | p-value: 0.7 beta: -0.10 |
| 21 | p-value: 0.0003, beta: -1.1 | p-value: 0.4, beta: -0.25 |
| 22 | p-value: 0.03, beta: -0.77 | p-value: 0.4, beta: -0.16 |
| 23 | p-value: 0.01, beta: -0.93 | p-value: 0.3, beta: -0.18 |

Table A2: Effect of varying threshold IOP on the linear associations between LC strains and retinal sensitivity.

According to this analysis, our conclusion still holds from the threshold IOP of 19 to 23. With 18 mmHg as a threshold value, our HTG subjects no longer show significant association between strains and IOP elevation. In general, the higher the IOP elevation, the stronger the negative correlation (beta values become more negative as threshold IOP is higher). It should be noted that for this experiment, the threshold IOP of 21 mmHg was still the most ideal in terms of splitting subjects equally between both groups.

F. **Adjusting Strains to the IOP-raised**

We adjusted our effective strain based on the level of elevated IOP (elevated IOP minus baseline IOP), by assuming a linear relationship between the two parameters. Since there are no known non-linear functions that approximate the relationship, and by Occam’s razor (the simplest explanation is usually the best one), we adjusted our measured strains with respect to the average level of IOP increase (18 mmHg), based on the following formula.

$$Adjusted Effective Strains=Measured Effective Strain \times\frac{18 mmHg}{Elevated IOP-Baseline IOP}$$

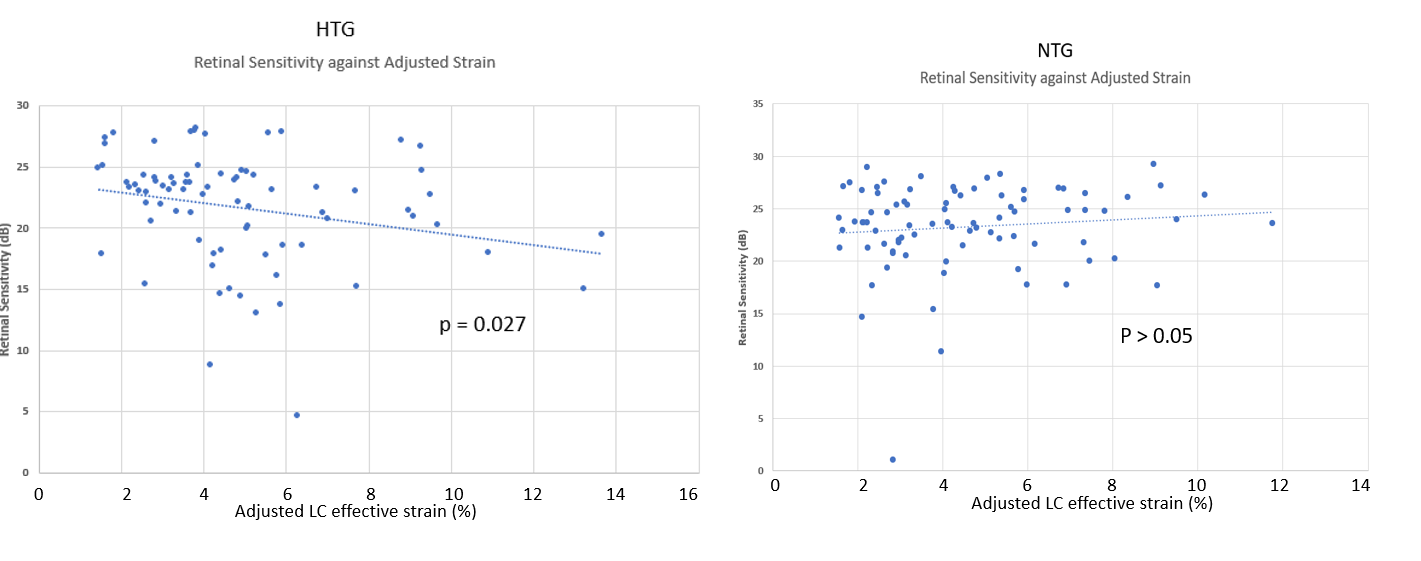

Figure A2: A scatterplot between the average retinal sensitivity against adjusted LC effective strain in HTG (left) and NTG subjects (right).

Based on the adjusted strains (**Figure A2**), we arrived at the same conclusion, that we found significant negative linear relationship between retinal sensitivity and measured strains in HTG but not in NTG.

G. **Regression of ONH Strains to other Visual Field Indexes**

First, we investigated the relationship between MD against average retinal sensitivity and we found that MD had a very strong correlation to the average retinal sensitivity (Figure A3a). Using MD and PSD to regress against average ONH strains, we still found a stark contrast in NTG and HTG subjects (Figure A3b-c). HTG subjects exhibited strong correlations between IOP-induced strains and MD/PSD values whereas such correlations were weak for NTG subjects. All conclusions stated in the manuscript remained unchanged when using MD values. When using PSD values, the correlations were only significant in HTG subjects for the LC tissue and not-significant (p>0.05) in other tissues.

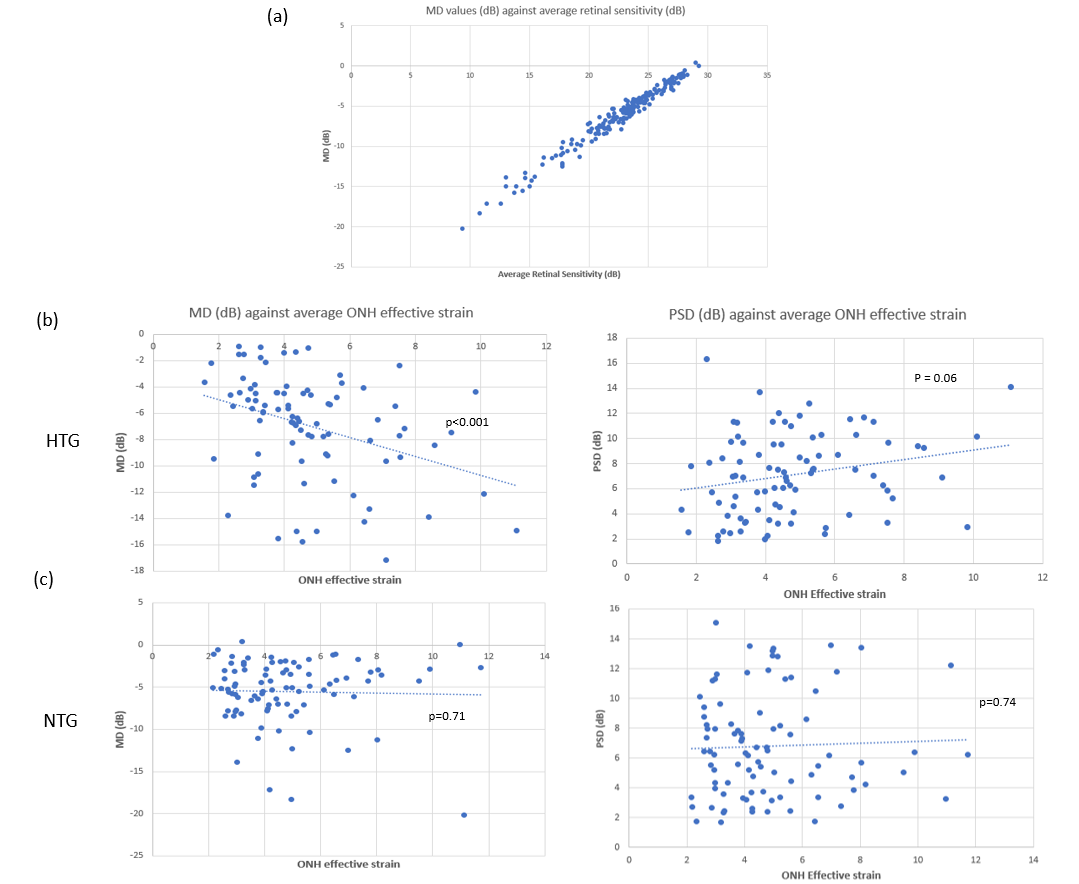

Figure A3: (a) Relationship between MD and average retinal sensitivity; (b) Relationship between MD and PSD against average ONH effective strain in HTG subjects; (c) Relationship between MD and PSD against average ONH effective strain in NTG subjects.

We have also performed the regional analysis with respect to TD and PD values. For HTG subjects, we still found significant negative correlations between regional TD/PD values and regional ONH effective strain (p-value<0.001 for TD, p = 0.03 for PD). For NTG subjects, we found negative correlations, but they were not significant (p-value = 0.06 for TD, p = 0.3 for PSD).

H. **Regression of ONH Strains to Regional RNFL thickness**

We found significant negative correlations between RNFL thickness and the corresponding average effective strain in each region (Superior, Inferior, Nasal and Temporal) for HTG subjects (Figure A4a). Thus, the regional strains were higher in the region with more local damage as indicated by thinning of the RNFL. This relationship was absent for NTG subjects (Figure A4b). This new analysis yielded results with similar implication to the one we presented in the manuscript.

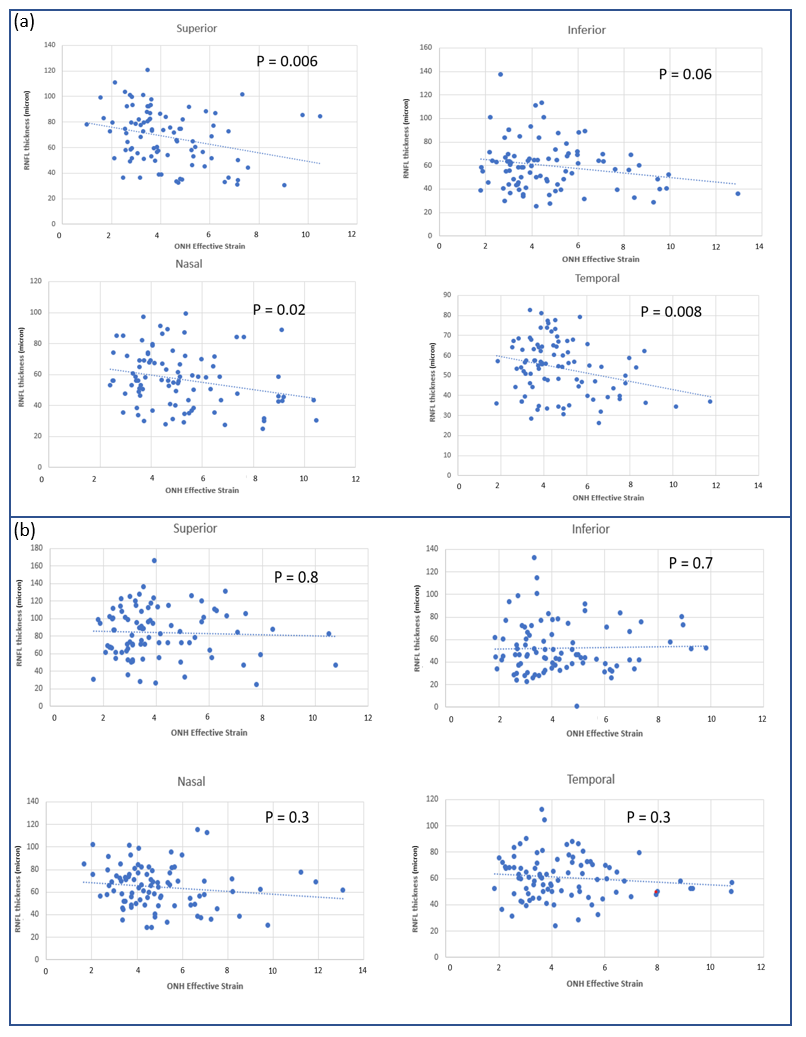

Figure A4: Relationship between regional RNFL thickness and the regional ONH effective strain for (a) HTG subjects (b) NTG subjects.

1. Downs, J. Crawford, Michael D. Roberts, and Claude F. Burgoyne. "The mechanical environment of the optic nerve head in glaucoma." *Optometry and vision science: official publication of the American Academy of Optometry* 85.6 (2008): 425.
2. Strouthidis, Nicholas G., and Michael JA Girard. "Altering the way the optic nerve head responds to intraocular pressure—a potential approach to glaucoma therapy." *Current opinion in pharmacology* 13.1 (2013): 83-89.
3. Sigal, Ian A., et al. "Eye-specific IOP-induced displacements and deformations of human lamina cribrosa." *Investigative ophthalmology & visual science* 55.1 (2014): 1-15.
4. Voorhees, Andrew P., et al. "Lamina cribrosa pore shape and size as predictors of neural tissue mechanical insult." *Investigative ophthalmology & visual science* 58.12 (2017): 5336-5346.
